# Characterization of Woodsmoke Generated in the Air Pollution Exposure Lab and Comparison to Diesel Exhaust

**DOI:** 10.1101/2025.02.27.640457

**Authors:** Yu Xi, Kristen Hardy, Vikram Choudhary, Julia Zaks, Carley Schwartz, Christopher F. Rider, Allan K. Bertram, Christopher Carlsten

## Abstract

To address the increasing concern of woodsmoke (WS) and better understand its effects on human health, a woodsmoke generation system was built in the Air Pollution Exposure Laboratory to facilitate future controlled human exposure studies. Two different woodsmoke conditions, flaming (WSFL) and smoldering (WSSM), were generated, and PM_2.5_ concentrations of approximately 500 μg/m^3^ were achieved. The woodsmoke produced using the system was characterized in this study and compared with diesel exhaust (DE) generated and collected at the same facility. Within the gas phase generated by the pollutants, WS showed slight increases in CO and CO_2_ compared to filtered air (FA), while DE contained significantly higher levels of NOx, CO_2_, and total volatile organic compounds compared to FA. The WS aerosols were comprised of approximately 98% organics, 0.6% NH_4_, 0.9% NO_3_, and 0.2% SO_4_. Among the organic species, the CHO1 and CHOgt1 families encompassed around 60%, which was higher than the fraction of oxygenated families in DE aerosols. Moreover, the WS aerosols had higher concentrations of Cd compared to the DE aerosols. Greater oxidative potentials were also observed for WSFL and WSSM compared to DE, with DTT consumption rates normalized to the PM mass being 0.0099 and 0.0090 nmol/min/μg, respectively. The difference in the compositions and properties of WS and DE suggests that it is critical to conduct further studies on how these pollutants can affect health differently.

## Introduction

Air pollution is a serious contributing factor to disease and premature death worldwide. In 2019, air pollution was attributable in approximately 6.6 million deaths globally (Murray et al., 2020). Air pollution was also reported to be the second-largest risk factor in global disease burden, measured in disability-adjusted life years (Murray et al., 2020). Fine particulate matter, characterized as particles less than 2.5 microns in diameter (PM_2.5_), is able to travel deep into the lungs when inhaled, circulate in the bloodstream, and cause systemic inflammation and oxidative stress (Xing et al., 2016). PM_2.5_ is one of the major constituents of air pollution that is known to cause and exacerbate a number of health conditions, including cardiovascular disease, a variety of cancers, and respiratory diseases, such as asthma and chronic obstructive pulmonary disease (COPD) (Xing et al., 2016).

Among the many types of air pollution, woodsmoke (WS) has been of increasing concern in recent decades due to the substantial increases in wildfires and associated health effects. WS is produced by the combustion of wood and is a complex mixture of gasses, particulate matter (including PM_2.5_), trace elements, hydrocarbons, free radicals, and more (Cascio, 2018a). Many of these compounds are proven to be detrimental to human health, with some being mild irritants or allergens and many being carcinogens (Naeher et al., 2007). One significant contributor to WS are forest fires, the number and severity of which has been steadily increasing in many areas of the world since the 1980s (Natural Resources Canada, 2023; United States Environmental Protection Agency, 2024). One study looking at the average outdoor PM_2.5_ levels over a 30-year period in the western United States found that during intense wildfires, 90% of the ambient particle pollution was attributable to the fires, and they were also the primary cause of the variances seen in summertime particle pollution levels from year to year (Xie et al., 2020). Additionally, domestic usage of wood as fuels is conducive to poor air quality. According to the World Health Organization, as of 2023, approximately 2.3 billion people relied on solid fuels such as wood to power their stoves, leading to indoor pollutant levels that were upwards of 100 times above the acceptable levels (World Health Organization, 2023). The use of wood in the home occurs all over the world; for example, in British Columbia, WS emissions from home heaters alone made up over 25% of PM_2.5_ in the ambient air (Government of British Columbia, 2023).

Given its considerable contribution to air pollution and potential detrimental health impacts, it is necessary to investigate the health effects of exposure to WS and the corresponding mechanisms by which it damages human health. Findings from *in vivo* and epidemiological studies have shown strong associations between exposure to WS and overall mortality, respiratory morbidity, as well as worsening of respiratory illnesses such as asthma and COPD (Cascio, 2018b). *In vitro* studies have further supported the link between WS and disease, with one study finding increases in DNA-damage, cell cycle disturbances, and interruptions to metabolic activities when several cell lines were exposed to WS particles (Hansson et al., 2023). Despite the large availability of literature exploring the effects of WS on human health, the number of studies which involve controlled human exposures to WS remains small. These types of studies aim to give a more precise view of how the human body is impacted by WS, as they can reduce confounding factors that other types of studies, namely animal and epidemiological studies, often suffer from. Two reviews of the available controlled human WS exposure studies were conducted in 2020 (Schwartz et al., 2020) and 2021 (Andersen et al., 2021), and identified 22 and 12 relevant studies, respectively. Both reviews focused on WS’s effects on lung function, airway inflammation, circulating cells and proteins, cardiovascular physiology, and oxidative stress, with one of the reviews additionally summarizing findings on systemic inflammation, thrombogenicity, and genotoxicity. The findings from both publications were in agreement with each other, with both stating that there were significant variations between the studies in emission source, exposure conditions, study design, and endpoints examined, making it difficult to compare findings (Andersen et al., 2021; Schwartz et al., 2020). Although both reviews concluded that no clear effect of WS on lung function has been found in the current human exposure studies, this was substantially due to the conflicting findings in combination with the vast differences in overall study designs.

The air pollution exposure laboratory (APEL) is equipped with an exposure booth allowing for controlled human exposure studies, where various studies about the effects of diesel exhaust (DE) on the respiratory system have been carried out (Carlsten et al., 2016; Jiang et al., 2014; Orach et al., 2023; Ryu et al., 2022; Thomson et al., 2021). A woodsmoke generator was built in APEL, modeled after a system developed at the United States Environmental Protection Agency, that can carry generated WS into the exposure booth, allowing for controlled human WS exposures. An understanding of the properties and composition of the WS produced is essential to ensure the safety of future participants, who would be exposed to this generated WS. Additionally, not only will having a deeper understanding of the physicochemical properties of the WS allow for more insight into the specific components which cause the physiological reactions observed in participants, but it will also aid in normalizing the exposure conditions to other studies, helping to create consistency within this relatively novel field of research. In the current study, the concentrations and properties of the PM_2.5_ generated by the combustion of lodgepole pine wood under smoldering and flaming conditions in APEL were investigated and compared with DE, the other air pollution type generated in APEL. Understanding how these different pollutants can affect health differently is necessary to address the evolving risk of air pollution and improve the health and safety of the communities.

## Experimental Methods

### Generation of WS

Dried lodgepole pine logs were obtained from the U.S Forest Service Missoula Fire Sciences, and were stored at ∼19°C in an airtight storage container. The wood was shaved using a block plane and then ground into uniform powders. This ‘sawdust’ was then dried in an oven for 90 minutes at 80°C in an aluminum pan. The dried wood was stored in a Falcon tube with desiccators to minimize moisture.

Figure 1 illustrates the APEL woodsmoke-generating system used for these experiments. The ground and dried lodgepole pine was aligned inside of a 96” quartz tube, and was heated by a ceramic heater, with the temperature controlled by a variable transformer. For flaming conditions, the variable transformer was set to 117W, producing measured temperatures in the ceramic heater of 649°C - 746°C (average 719°C). For smoldering conditions, the variable transformer was set to 65W, producing measured temperatures in the ceramic heater of 278°C - 340°C (average 319°C). A track controlled by a servo-motor moves the ceramic heater along the tube at a consistent slow rate, allowing for continuous burning of fresh wood. On one end of the quartz tube is a source of combustion air which pumps air at an adjustable flow rate into the tube; on the other end is a Pyrex piece, which attaches to a system of piping which leads into the exposure booth. Near this Pyrex piece is a supply of dilution air, the flow rate of which is also adjustable. The WS from the combustion mixes with the dilution air in the Pyrex fitting, and this mixture is pumped into the exposure system, which was described in detail by Birger *et al*. (Birger et al., 2011).

**Figure 1.**
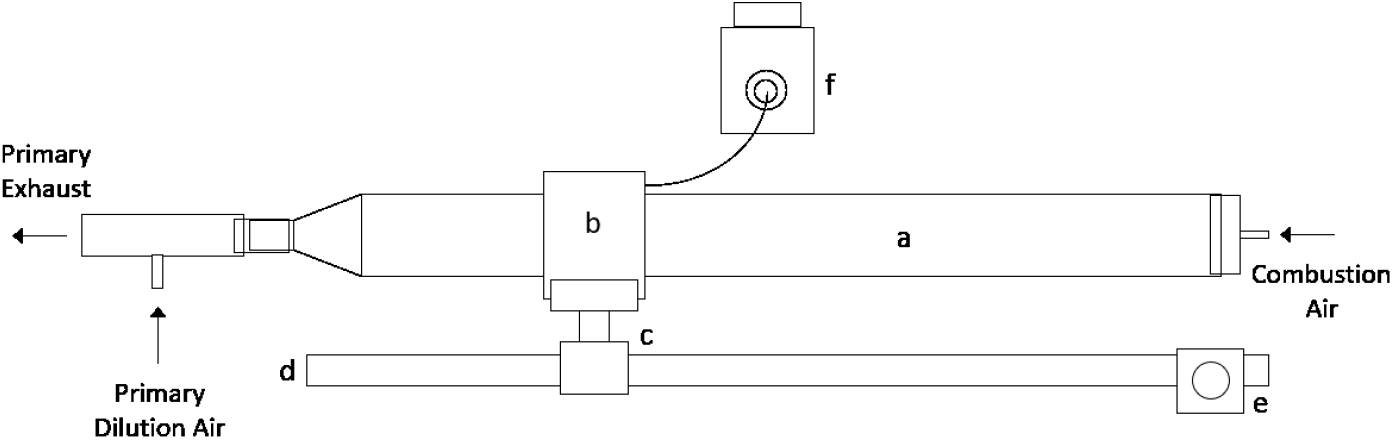
Schematic of the woodsmoke generator. The parts of the generation system: a) Quartz tube; b) Ceramic heater; c) Custom heater stand; d) Furnace track; e) Track servo-motor; f) Variable transformer.

### In-booth Woodsmoke Characterization

To understand the WS generated in this study, the concentrations of PM_2.5_ and gasses in the exposure booth were monitored, and in-booth samples were collected for further characterization. PM_2.5_ concentrations were measured using a Rupprecht & Pattashnick 1400a Tapered Element Oscillating Microbalance (TEOM) monitor, with size distributions detected using a TSI Model 3936 Scanning Mobility Particle Sizer (SMPS). The chemical composition of the aerosols generated during the different conditions were tested using a time-of-flight Aerosol Mass Spectrometer (ToF AMS; Aerodyne).

PM_2.5_ was collected on several types of filters and grids for different analyses. The collection was conducted using a Harvard aerosol impactor with a PM_2.5_ inlet, which was connected to the exposure booth during the exposure runs with a flow rate of 10 L/min. PM_2.5_ samples were collected on carbon-coated transmission electron microscopy (TEM) grids and imaged with a FEI Tecnai G2 Twin TEM operated at an accelerating voltage of 200 kV. Sample collections on 37mm Pallflex tissue quartz filters were analyzed for elemental carbon/organic carbon (EC/OC) ratios with the NIOSH5040 method using a Sunset Laboratory OCEC Analyzer M5L. Elemental analysis was conducted with samples collected on Environmental Express 37mm PTFE filters using an Agilent 7700x ICP-MS after leaching with aqua regia and rinsing with deionized water. WS samples collected on 37mm quartz fiber filters were used for the endotoxin and oxidative potential analyses.

### Endotoxin Analysis

The endotoxin concentrations of WS samples were analyzed using the gel-clot assay with the Charles River Endosafe Limulus Amebocyte Lysate (LAL, detection limit: 0.06 EU/mL). The Charles River Endosafe Control Standard Endotoxin (CSE) and Limulus amebocyte lysate reagent water (LRW) were used as the positive and negative controls, respectively. All glassware and filters were depyrogenated at 250°C for 2 hours, then allowed to cool prior to collection and testing. The samples were extracted from the filters with LRW via 2-hour shaking at 600 rpm, and the supernatants of the extraction were used in the endotoxin tests. The tests were conducted by combining 0.1 mL of LAL with 0.1 mL of the samples or the controls in 10 × 75 mm glass tubes and incubating the mixtures at 37±1 °C for 60 ±1 minutes. The presence of firm gel after the incubation served as an indicator of the presence of endotoxin at the detection limit.

### Oxidative Potential Test

An assay using PM to catalyze the reduction of oxygen by dithiothreitol (DTT) was developed by Cho *et al*. (Cho et al., 2005), and was used to determine the oxidative potential of the PM. The PM_2.5_ was extracted from filters using 0.1M phosphate buffer, followed by 2-hour shaking at 600 rpm. A total of 250μL of each filter extract, as well as 250μL of 0.1M phosphate buffer (serving as a blank), were incubated at 37°C, each with an addition of 250μL of 200μM DTT in phosphate buffer, for time intervals ranging from 5 to 40 minutes. The reactions were stopped using trichloroacetic acid at the designated times, then DTNB and Tris-Base were added, and the absorbance of the solutions was measured at 412nm using a SpectraMax® i3x plate reader. The absorbance readings were compared against a standard curve of DTT in phosphate buffer to calculate the concentrations of DTT.

## Results and Discussions

### Particle and gas metrics

The concentrations and sizes of particles generated during the DE, FA, WSFL, and WSSM conditions are listed in Table 1. As expected, both the mass and particle concentrations of the pollution conditions were significantly higher than that of the FA control condition, with the target PM_2.5_ concentration being 300 μg/m^3^ for DE and 500 μg/m^3^ for WS to be consistent with previous studies (Ghio et al., 2012; Rebuli et al., 2019; Volpi et al., 2019). Due to the unpredictable and inconsistent composition of the wood particles in comparison to the diesel, maintaining the target PM_2.5_ concentrations was more difficult for WS than for DE, as shown by the higher standard deviations seen with both WSFL and WSSM. The median particle sizes did not vary significantly between the DE, WSSM, and WSFL conditions, with all being around 100 nm. This is in agreement with the size distributions shown in Figure 2, where peaks around 100 nm were observed for DE, WSSM, and WSFL conditions, while no clear peak was observed for the FA condition. Also, of note in Figure 2 is that the WSFL condition, compared to the WSSM condition, tended to have a higher proportion of particles smaller than 200 nm, and fewer particles >200 nm. This may explain the higher particle number concentrations observed for WSFL compared to WSSM under similar mass concentrations.

**Table 1.**
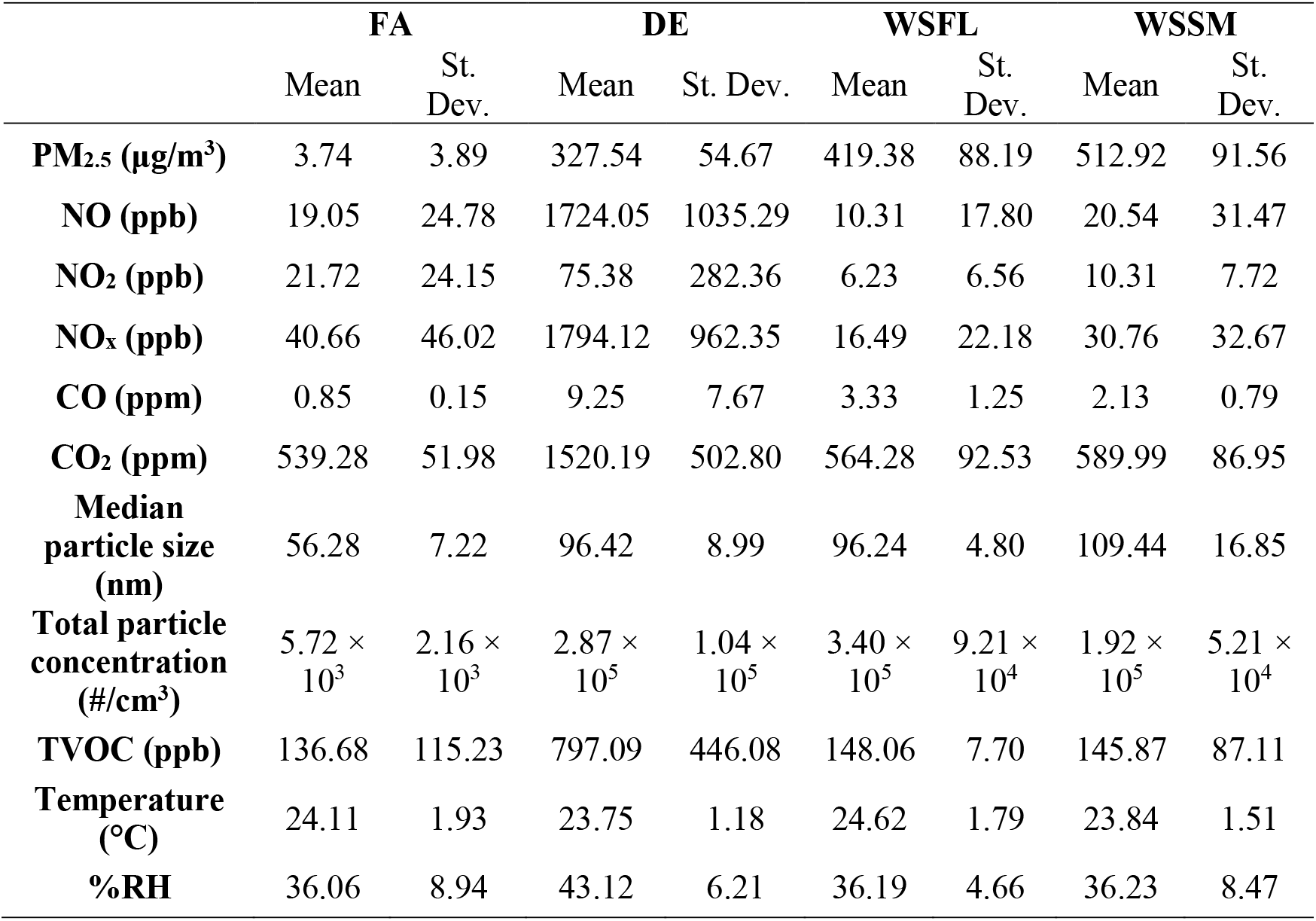
In-booth characteristics for filtered air (FA), diesel exhaust (DE), woodsmoke flaming (WSFL), and woodsmoke smoldering (WSSM). Means and standard deviations were calculated based on six duplicate runs, with the PM_2.5_ concentrations aimed at achieving nominal concentrations of 300 μg/m^3^ for DE and 500 μg/m^3^ for WS.

**Figure 2.**
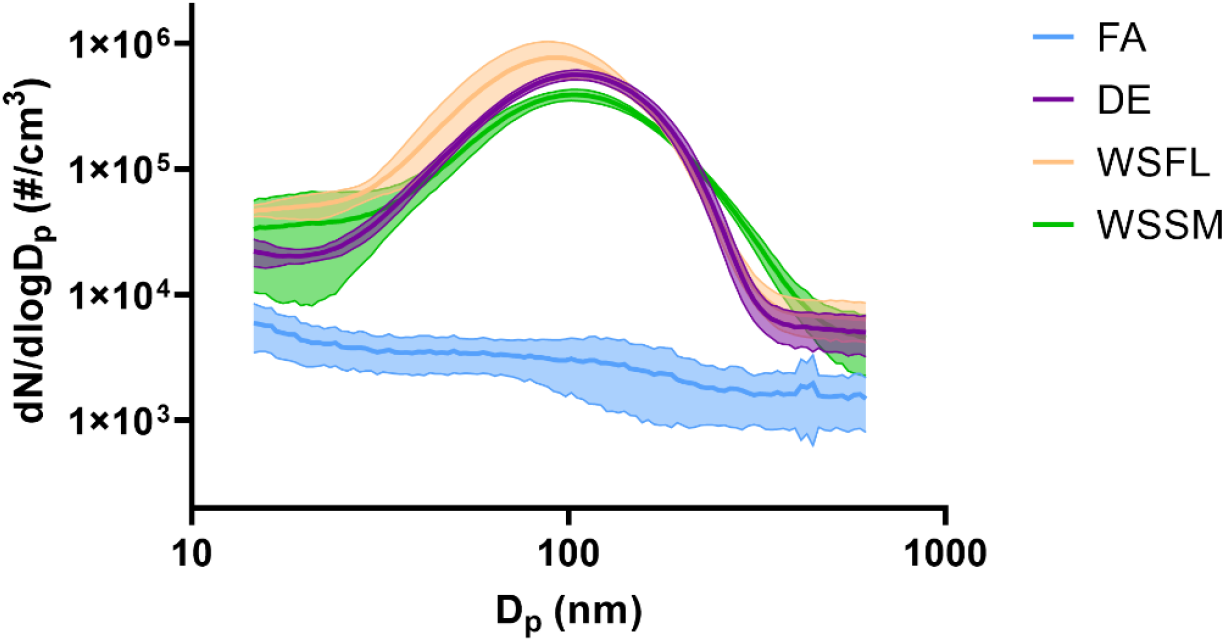
Size distributions of particles generated from DE, FA, WSFL, and WSSM conditions, error bands representing the standard deviations calculated based on four replicates.

Also included in Table 1 are the gas concentrations measured during the different conditions. DE contained the highest concentration of NO_x_, with an average of 1794.12 ppb NO and 75.38 ppb NO_2_; this is a stark difference from the WS conditions, where the concentrations of NO_x_ were similar to those seen in the FA condition. Production of CO was also observed during the diesel engine combustion, with an average CO level of 9.25 ppb being reached. Though not as high as that of DE, CO concentrations around 2.13 and 2.33 ppb were observed for WSSM and WSFL, respectively, which is still higher than the 0.85 ppb CO concentration observed for FA. The DE condition produced a higher concentration of CO_2_ as well, while the CO_2_ levels in the WS conditions were not significantly higher than that of the FA condition. Moreover, the total volatile organic compound (TVOC) measurements showed notably higher concentrations of TVOCs in the DE condition, while the TVOC concentrations of the WSFL and WSSM conditions were similar to that of FA. Higher concentrations of NO_x_, CO, CO_2_, and TVOC were present under the DE condition even at lower PM concentrations compared to the WS conditions.

The ratios of elemental carbon (EC) to organic carbon (OC) in the pollutants were measured for the aerosol samples collected during the DE, WSFL, and WSSM conditions (Table 2). A dramatic difference was seen between the DE samples, which had a mean EC/OC ratio of approximately 0.31, and the two WS samples, which contained minimal EC, and had EC/OC ratios below 0.01. The higher EC/OC ratio in DE aerosols is unsurprising, as EC mainly originates from combustion of fossil fuels. The low EC in WS aerosols was also seen in a previous study (Singh et al., 2023).

**Table 2.** Ratios of elemental carbon (EC) to organic carbon (OC) for aerosols collected during the DE, WSFL, and WSSM exposure conditions. Means and standard deviations were calculated based on four replicate runs for DE and three replicates each for WSFL and WSSM.

|  | EC/OC ratio |  |
| --- | --- | --- |
|  | Mean | St. Dev. |
| DE | 0.31 | 0.04 |
| WSFL | 0.0077 | 0.0026 |
| WSSM | 0.0075 | 0.0006 |

### Endotoxin Test

Endotoxin is a lipopolysaccharide found in the membranes of Gram-negative bacteria, and is released into the air upon bacterial cell lysis (Bertics et al., 2006). Endotoxins are present in wood and can be released into the air during wood burning leading to high exposure levels (Semple et al., 2010). Endotoxins are known to have pro-inflammatory effects on the respiratory tract, and have been implicated in the development of diseases such as asthma, organic dust toxic syndrome, and COPD (Bertics et al., 2006; Douwes et al., 2003; Liebers et al., 2008). Therefore, it is imperative to examine the endotoxin concentrations in the booth to ensure the safety of future participants undergoing controlled WS exposures.

The results of the LAL tests performed on all the PM_2.5_ sample extractions collected during the FA, DE, WSFL, and WSSM conditions were negative, suggesting that endotoxin concentrations in the booth were lower than 0.5 EU/m^3^ for all the four conditions. The endotoxin concentrations of the four conditions are similar to the reported values of endotoxin concentration in urban ambient air, which has been recorded to have a mean or median concentration between 0.006 and 5.7 EU/m^3^ (Rolph et al., 2018). This indicates that neither the DE or WS conditions created in the lab generated an endotoxin concentration significantly higher than that of urban ambient air. Additionally, the endotoxin levels produced during the exposures were lower than the recommended exposure limits set by the Health Council of the Netherlands, who recommend a concentration no greater than 90 EU/m^3^ for workers over an 8-hour time span (Health Council of the Netherlands, 2010). Thus, the endotoxin levels of the FA, DE, WSFL, and WSSM conditions can be confidently considered safe for human exposure.

### TEM Imaging

TEM images of particles generated during the different conditions are shown in Figure 3. Two main types of particles were found in all four conditions: soot aggregations and organic particles (Li et al., 2016). For the soot aggregations generated by DE (Figure 3a), WSFL (Figure 3e, 3f), and WSSM (Figure 3i, 3g), the diameters mostly lay in the range of 100 nm to 200 nm. The soot aggregate structures observed under both WS conditions had softer edges compared to those found in DE aerosols, with one possible reason for this difference being that the soot particles generated under the WS conditions were coated in organic matter.

**Figure 3.**
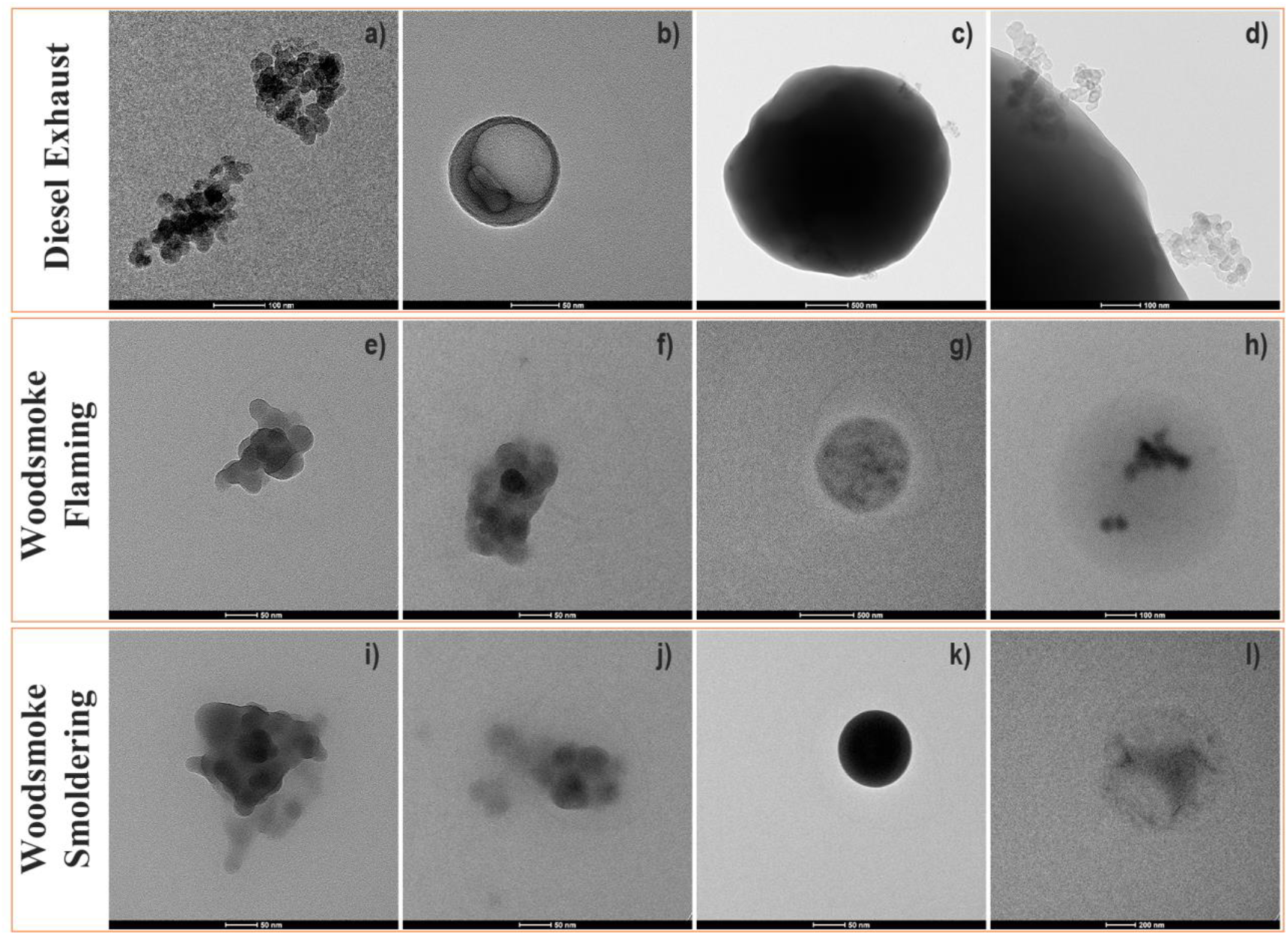
TEM images of PM collected from the booth during DE (a, b, c, d), WSFL (e, f, g, h), and WSSM (i, j, k, l) exposure conditions.

Organic particles found in the DE (Figure 3b, 3c), WSFL (Figure 3g, 3h), and WSSM (Figure 3k, 3l) conditions had a larger size range compared to the soot aggregates, with diameters ranging from 100 nm up to 2 μm. Moreover, an agglomerate structure of organic particles and soot was observed in DE aerosols (Figure 3c, 3d), and a core-shell structure was observed in WSFL aerosols (Figure 3h). One thing to consider is that organics are more fragile under the high vacuum and electron beams used for the TEM imaging and can be destroyed during the imaging process(Ault & Axson, 2017). This could explain the irregular shape seen in Figure 3l. Therefore, one limitation of TEM imaging in this study is that some of the structural and physicochemical characteristics of the PM_2.5_ may have been compromised during the imaging process.

## Metallic Compositions

The metallic compositions of aerosols generated during DE, WSFL, and WSSM exposures were measured using ICP-MS, and the mass fractions after subtracting out the blanks are shown in Figure 4. The metallic concentrations measured in the samples were compared against the blank filter concentrations before subtraction, and the corresponding p-values were calculated (Figure 4). Metal elements showing significant differences from the blank were found to be Na, P, Mn, Fe, Zn, Rb, Mo, and Ag in DE aerosols, Na, Ag, and Cd in WSFL aerosols, and Na, K, Rb, Mo, Ag, Cd, and Pb in WSSM aerosols. Metals including P, Zn, and Mo were found to be present in statistically significant concentrations compared to the blank in DE, but not in either WS condition.

**Figure 4.**
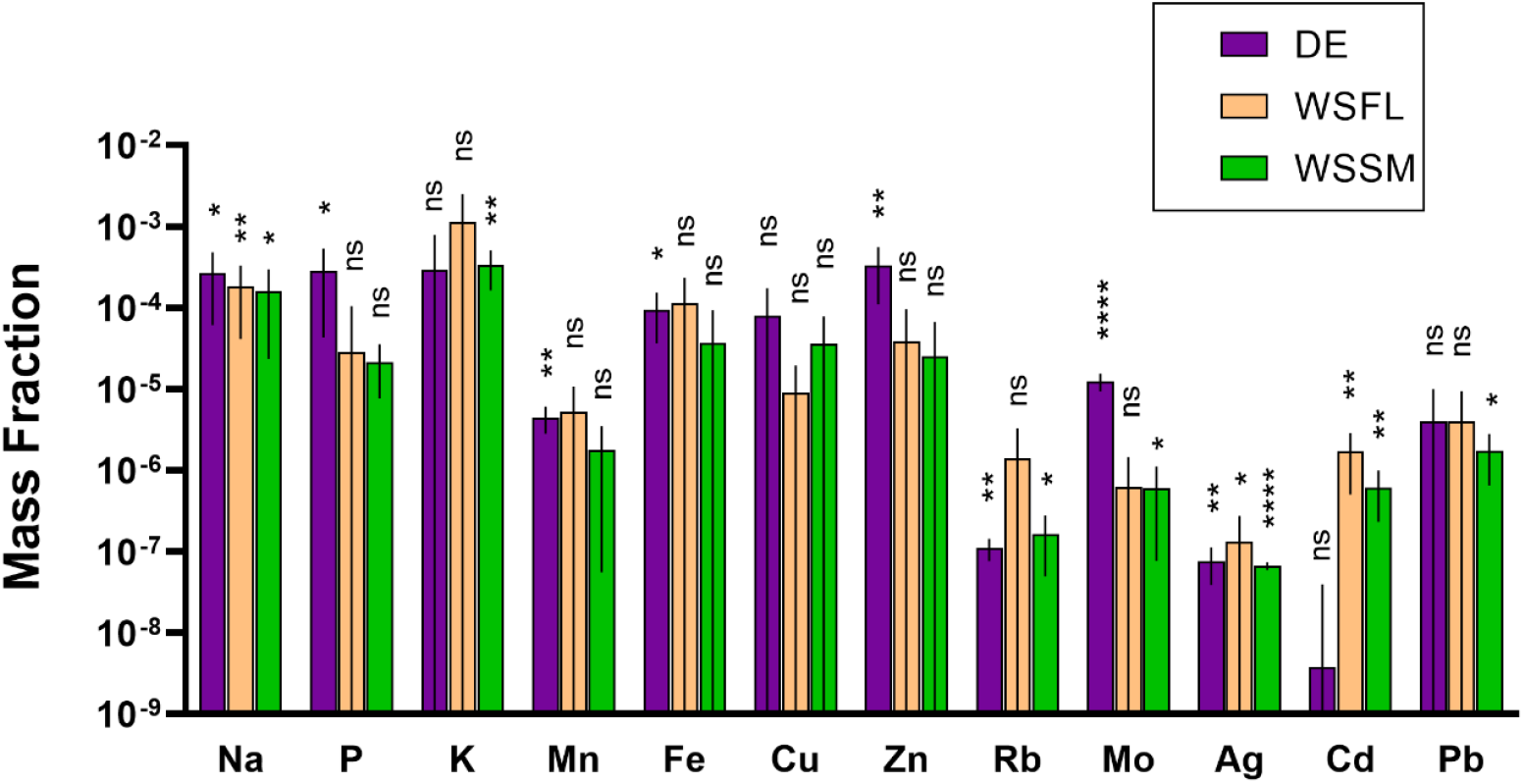
Mass fractions of different elements measured by inductively coupled plasma mass spectrometry (ICP-MS) collected from the booth on filters during DE, WSFL, and WSSM exposures. Error bars represent the SEM of three replicate tests, and p-values were calculated based on the comparisons between the samples and the blanks.

Most notable from these results was the discrepancy in the mass fractions of Cd; statistically significant fractions of Cd were found in the WSFL and WSSM aerosols, but not in the DE aerosols. With Cd playing a role in the development and/or exacerbation of numerous respiratory diseases including emphysema, asthma, and bronchitis, the fractional difference noted between the two pollutants offers a new avenue of potential investigation in future pollution toxicology studies (Charkiewicz et al., 2023).

### Chemical Compositions Measured by AMS

The chemical composition of the aerosols in the DE, WSFL, and WSSM conditions measured by the AMS are shown in Figure 5. For all three conditions, organic components took up the majority of the aerosol fractions (over 98%), with DE aerosols containing a slightly higher mass fraction of organic components compared to the two WS conditions (Figure 5a). The mass fractions of chemical families within the organic species showed a significant difference between DE and WS conditions (Figure 5b). Around 85% of the organic species in DE aerosols belong to the CH family, while in the WSFL and WSSM aerosols, the CH family constitutes only around 33% of the organic species present. Conversely, the CHO1 and CHOgt1 families were seen in significantly greater proportions in WSFL and WSSM aerosols compared to in the DE aerosols. The difference shows that there are more oxygenated organic species generated under WS conditions than under DE conditions. The same conclusion can also be drawn from the comparison of O:C ratios, where the O:C ratios were seen to be approximately 0.07, 0.54, and 0.58 for DE, WSFL, WSSM, respectively (Figure 5c). Different organic families showed similar mass fractions for the aerosols generated under the WSFL and WSSM conditions (Figure 5b), with the O:C ratio for WSSM slightly higher than that of the WSFL (Figure 5c).

**Figure 5.**
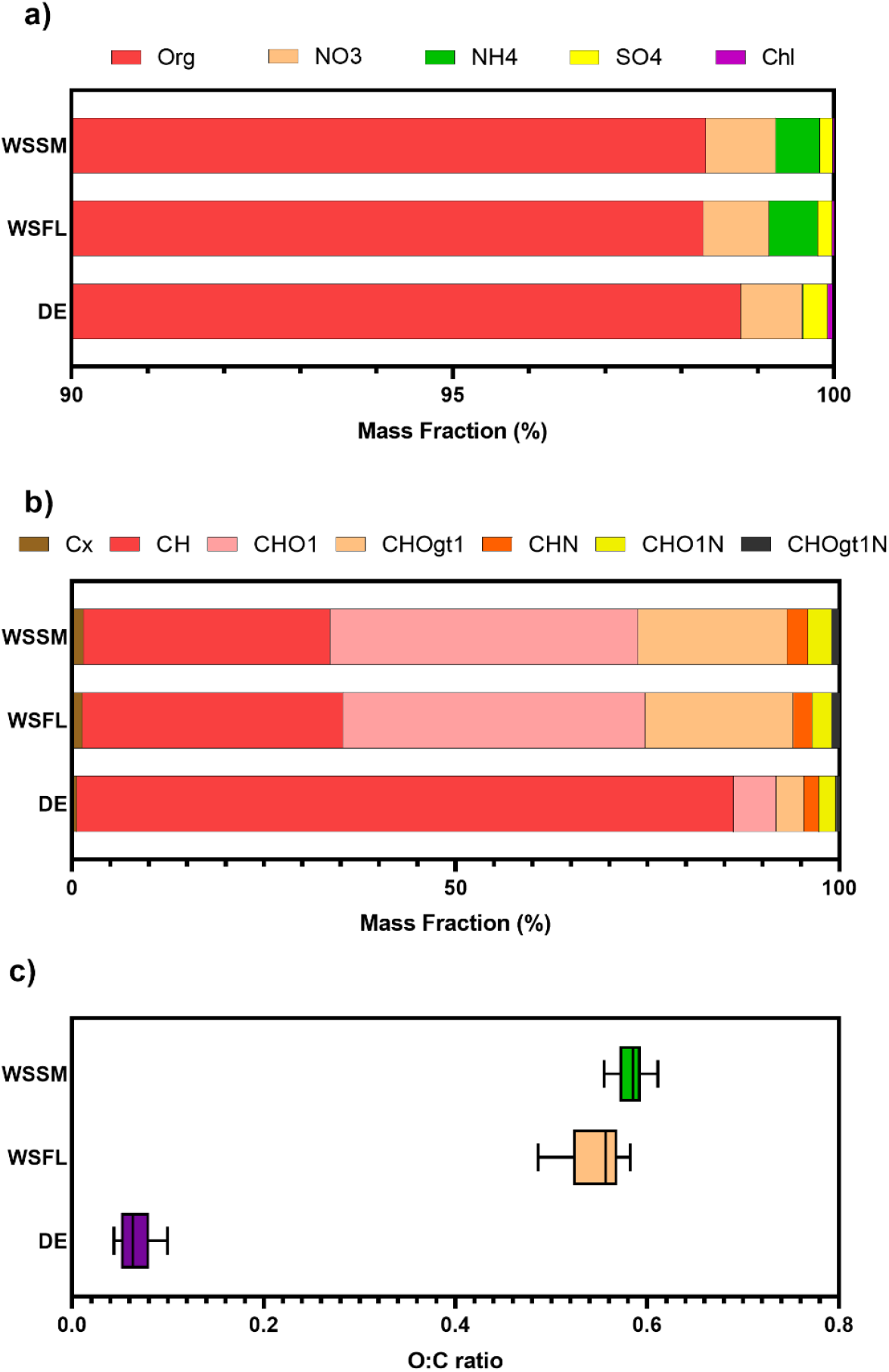
Mass fractions of relevant chemical families in a) overall species and b) organic compositions for the aerosols produced under DE, WSFL, and WSSM conditions. c) O:C ratio for the aerosols sampled for the DE, WSFL, and WSSM conditions, with error bars showing the standard deviations.

**Figure 5.**
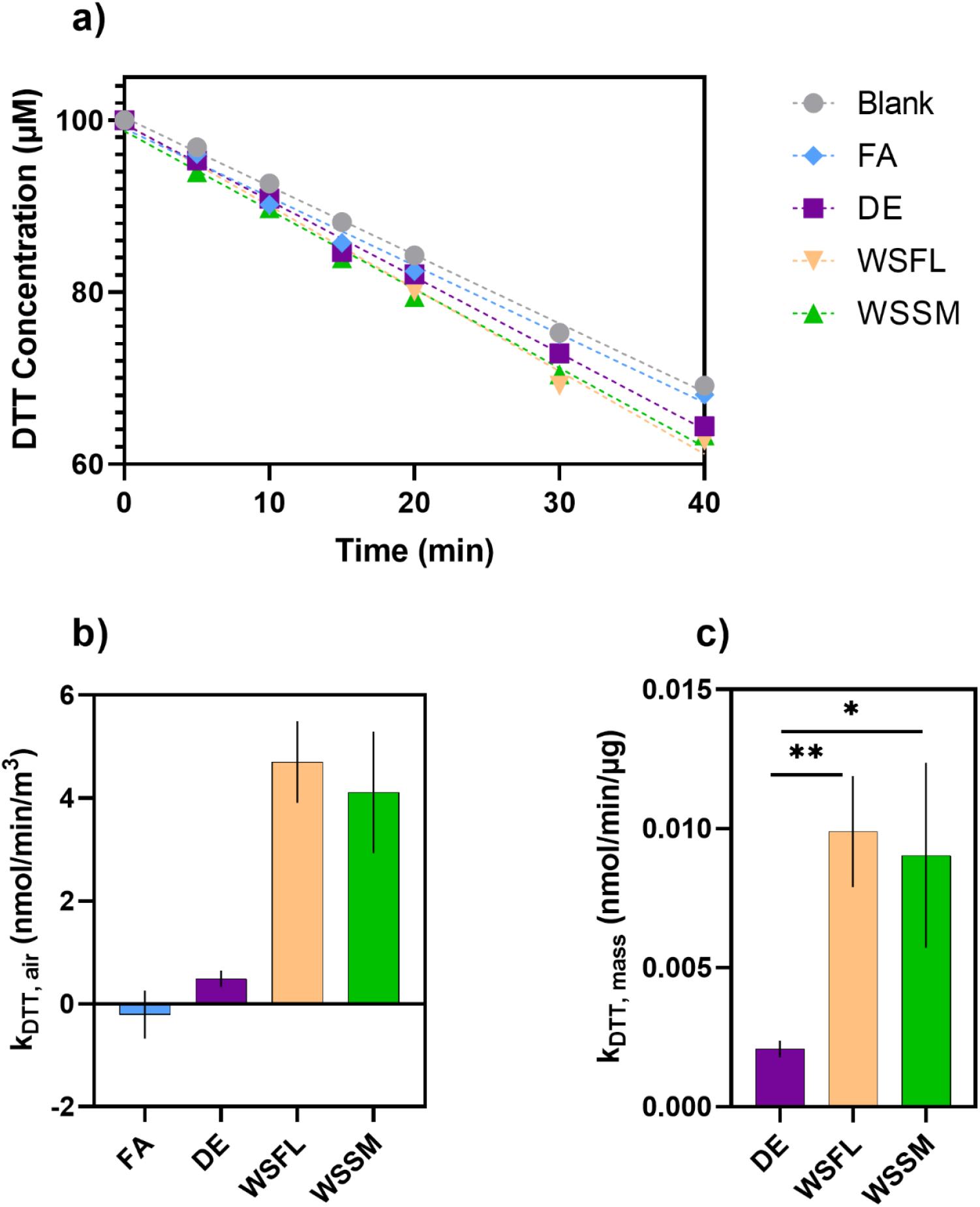
Oxidative potential measured by the DTT assay. a) Change in DTT concentration over time with the addition of water extractions from sample filters collected from the booth during DE, WSFL, WSSM, and FA exposures, compared to the blank. b) Rate of DTT loss normalized to volume of air for DE, WSFL, WSSM, and FA samples, with error bars representing standard deviations of three replicates. c) Rate of DTT loss normalized to PM mass of DE, WSFL, and WSSM, with error bars representing standard deviations of three duplicates.

Other components present in all the three conditions included SO_4_, which is mostly sourced from sulfate salts, and NO_3_, which is potentially sourced from nitrate salts or organic nitrogen. The DE aerosols contained slightly higher fractions of SO_4_ (0.32%) than the WSFL and WSSM aerosols (0.19% and 0.17%, respectively). NO_3_ comprised about 0.81%, 0.86%, and 0.92% of the organic species for the aerosols generated under DE, WSFL, and WSSM conditions, respectively. Additionally, both the WSFL and WSSM aerosols contained approximately 0.6% NH_4_, while the DE aerosols contained negligible amounts (Figure 5a). The NH_4_ seen in the WS conditions can be attributed to ammonium salts, which can form via the reaction of NH_3_. NH3 has been shown to be produced during biomass burning, and thus the presence of NH4 in the WS conditions aligns with previous studies (Bouwman et al., 1997; Feng et al., 2023).

### Oxidative Potential

Studies have shown that the adverse health effects of PM can be partially attributed to oxidative stress, which can be caused by reactive oxygen species (ROS) generated by PM (Valavanidis et al., 2013; Weichenthal et al., 2016; Yang & Omaye, 2009). The ability to generate ROS is called oxidative potential (OP). In this study, the OP of the water extractions of PM samples was measured using a DTT assay. Figure 5a illustrates the change in DTT concentrations for different samples in one trial, showing that the DE, WSFL, and WSSM samples were able to accelerate the oxidation of DTT compared to the blank. As expected, the reaction rate of DTT with the FA sample extract was similar to that of the blank. The reaction rates were calculated by subtracting the rate of DTT loss of the blank and then normalized to the volume of air (k_DTT, air_, Figure 5b) or mass of PM (k_DTT, mass_, Figure 5c). The average k_DTT, mass_ for DE, WSFL, and WSSM samples were 0.0021, 0.0099, 0.0090 nmol/min/μg, respectively. The oxidative potential of the DE sample was significantly lower than that of the WS samples, while no significant difference was found between the oxidative potentials of the two WS conditions. One possible reason for the difference between the oxidative potentials of DE and WS samples is that the WS aerosols contain more oxygenated species, as shown in the chemical compositions tested by AMS.

Previous studies measuring OP for aerosol samples collected in Greece and California during wildfire events observed k_DTT, air_ around 0.2 – 0.3 nmol/min/m^3^ and k_DTT, mass_ around 0.03 – 0.04 nmol/min/μg (Fang et al., 2023; Mylonaki et al., 2024). The OP of the WS aerosols generated in the current study were higher than the wildfire aerosols when normalized to the volume of air, but lower when normalized to the mass of PM. The higher OP per volume of air of the in-lab WS aerosols can be explained by higher pollution concentrations due to negligible transport distance and lower dilution rate compared to the in-field samples. On the contrary, the complexity of the wildfires, along with the possible oxidation during the transportation of the WS, can contribute to the higher OP per mass of PM compared to the WS generated in the current study.

## Conclusions

In the current study, a woodsmoke generation system for controlled human exposure studies was designed, and the WS generated under flaming and smoldering conditions was characterized. With the PM_2.5_ concentration of WS aimed at 500 μg/m^3^, slight increases in CO_2_ and CO were observed compared to FA controls, and no generation of NO_x_ was found. The levels of TVOC concentrations for the WSFL and WSSM conditions were also similar to those of the FA controls. The median particle sizes for the WSFL and WSSM conditions were all found to be approximately 100 nm. The EC/OC test showed that most of the carbon in the two WS conditions was organic. The endotoxin concentrations in the WS aerosols were below the detection limit of 0.5 EU/m^3^. TEM imaging showed the existence of soot and organic particles. Na, Ag, and Cd were observed in the WSFL aerosols, and Na, K, Rb, Mo, Ag, Cd, and Pb were observed in the WSSM aerosols. From the measurement of the AMS, more than 98% of the WS aerosols were composed of organics, with NH_4_, NO_3_, and SO_4_ representing 0.6%, 0.9%, and 0.2% of the aerosols, respectively. Within the organic chemicals for WS aerosols, the CH family comprised around 33% of the organic mass fraction, and the CHO1 along with the CHOgt1 comprised around 60%. The oxidative potentials of the WSFL and WSSM water extractions measured by k_DTT, mass_ were 0.0099 and 0.0090 nmol/min/μg, respectively.

The generated WSFL and WSSM showed similar physicochemical properties in general. The WSFL conditions tended to contain larger proportions of smaller particles compared to the WSSM conditions, resulting in higher number concentrations of particles at similar mass concentrations. The aerosols generated in APEL under the two WS conditions were also compared with the DE condition. Some major differences found include: 1) DE conditions contained higher concentrations of NOx, CO_2_, and TVOC; 2) DE aerosols had higher EC/OC ratios; 3) WSFL and WSSM aerosols had higher fractions of Cd; 4) WSFL and WSSM aerosols had higher fractions of ammonium and oxygenated organic species; and 5) WSFL and WSSM aerosols extracted in water had higher oxidative potentials. Given the higher fractions of Cd, oxygenated organic species and oxidative potential in WS, controlled human exposure studies are necessary to enhance the understanding on how WS can affect respiratory health in comparison to DE.

